# Noise in a metabolic pathway leads to persister formation in *Mycobacterium tuberculosis*

**DOI:** 10.1101/2021.11.26.470138

**Authors:** Jeffrey Quigley, Kim Lewis

**Author notes:** Corresponding author: Kim Lewis.

## Abstract

Tuberculosis is difficult to treat due to dormant cells in hypoxic granulomas, and stochastically-formed persisters tolerant of antibiotics. Bactericidal antibiotics kill by corrupting their energy-dependent targets. We reasoned that noise in the expression of an energy-generating component will produce rare persister cells. In sorted low ATP *M. tuberculosis* grown on acetate there is considerable cell-to-cell variation in the level of mRNA coding for AckA, the acetate kinase. Quenching the noise by overexpressing *ackA* sharply decreases persisters, showing that it acts as the main persister gene under these conditions. This demonstrates that a low energy mechanism is responsible for the formation of *M. tuberculosis* persisters and suggests that the mechanism of their antibiotic tolerance is similar to that of dormant cells in a granuloma. Entrance into a low energy state driven by stochastic variation in expression of energy-producing enzymes is likely a general mechanism by which bacteria produce persisters.

## Introduction

Tuberculosis is the most important infectious disease caused by a bacterial pathogen and is responsible for killing 1.4 million people a year (WHO, 2019). This ongoing global epidemic stems from the difficulty of eradicating the pathogen with currently available antibiotics. Treatment of antibiotic-susceptible *M. tuberculosis* requires 6 months with a combination of rifampicin, isoniazid, ethambutol, and pyrazinamide (Zumla, Nahid, & Cole, 2013). Not surprisingly, this results in side effects and poor compliance. The need for a lengthy treatment is attributed to the presence of dormant, non-replicating *M. tuberculosis* cells that are tolerant of killing by antibiotics (Datta et al., 2016; Sacchettini, Rubin, & Freundlich, 2008; Sonnenkalb et al., 2021).

The disease is typically associated with the walling off of *M. tuberculosis* in a hypoxic, acidified granuloma, a complex structure made primarily of immune cells and their products (Guirado & Schlesinger, 2013). These various stressors induce a population-wide low metabolic state termed dormancy. DosR and PhoP regulators control entrance of cells into a non-replicative state under hypoxia and acid stress (Baker, Johnson, & Abramovitch, 2014; Leistikow et al., 2010; Namugenyi, Aagesen, Elliott, & Tischler, 2017; Zheng et al., 2017), and RelA mediates starvation-induced non-replication (Dutta et al., 2019; Primm et al., 2000). Regardless of the stress, induction of dormancy is characterized by a metabolic downshift in the population and increased antibiotic tolerance (Boldrin, Provvedi, Cioetto Mazzabo, Segafreddo, & Manganelli, 2020; Gengenbacher, Rao, Pethe, & Dick, 2010; Wayne & Hayes, 1996). *M. tuberculosis* incapable of this metabolic downshift are more susceptible to antibiotics *in vivo* (Baek, Li, & Sassetti, 2011).

Apart from this population-wide response to distinct stress factors, *M. tuberculosis* also forms a small subpopulation of persister cells that are produced stochastically during normal growth and are tolerant of killing by antibiotics (Jain et al., 2016; I. Keren, Minami, Rubin, & Lewis, 2011; Srinivas, Arrieta-Ortiz, Kaur, Peterson, & Baliga, 2020; Torrey, Keren, Via, Lee, & Lewis, 2016). Persisters were originally discovered by Joseph Bigger in a population of *S. aureus* in 1944 (Bigger, 1944), and decades later they are attracting increased interest due to their role in recalcitrance of chronic diseases to antibiotic therapy (Lewis & Manuse, 2019). Tolerance is likely based on a shared feature of bactericidal antibiotics - killing by corrupting their targets (Iris Keren, Kaldalu, Spoering, Wang, & Lewis, 2004). For example, aminoglycosides such as streptomycin cause mistranslation, which leads to the production of toxic misfolded peptides (Davis, Chen, & Tai, 1986). Based on this, we suggested that persisters are low-energy cells (Brian P. Conlon et al., 2016). Indeed, persisters in *S. aureus* and *E. coli* have low levels of ATP (Brian P. Conlon et al., 2016; Manuse et al., 2021; Zalis et al., 2019). Sorting of cells treated with antibiotics with low levels of expression of TCA cycle enzymes enriches in persisters in these two species. However, the relative input of these enzymes into persister formation is unknown and detecting both expression of an energy producing component and ATP in the same cell has not been achieved yet. The mechanism by which multi-drug tolerant *M. tuberculosis* persisters are formed is largely unknown. Given that both population-wide dormancy of cells, and persisters that can form during growth exhibit antibiotic tolerance, understanding the mechanism of *M. tuberculosis* persister formation is critical for developing more effective therapies to treat this important disease.

Here, using single cell analysis, we demonstrate that *M. tuberculosis* persisters are stochastically generated low ATP cells. It is this low energy state that renders them tolerant of antibiotics. Further, using direct measurement of ATP and transcription noise in metabolic enzymes in the same cell, we explore the mechanistic basis of persister formation in *M. tuberculosis*. We show that, in a simple growth medium with acetate, low ATP cells express low levels of the acetate kinase AckA. By quenching noise through overexpression of AckA we are able to dramatically reduce the level of persisters, indicating that AckA functions as the main persister gene under these conditions. The approaches described in this study provide a means to determine the relative contribution of any gene into persister formation. Stochastic entrance into a low ATP state is likely a general mechanism of persister formation in bacteria.

## Results

### Low levels of ATP are linked to antibiotic tolerance in *M. tuberculosis*

In order to examine a causal link between a low energy state and persister formation in *M. tuberculosis*, we took advantage of the antibiotic bedaquiline that specifically inhibits the mycobacterial F1F0 ATP synthase by binding to its C-subunit (Andries et al., 2005). Adding bedaquiline to a growing culture reduced the level of ATP 2-fold after a relatively short, 4 hour exposure, as detected with luciferase (Figure 1A). In order to measure the level of persisters in these cells, cultures were washed to remove bedaquiline, challenged with either rifampicin + streptomycin (Rif/Strep) or isoniazid (INH) for 7 days, and viability was determined by colony count. Pre-treatment with bedaquiline significantly increased the number of antibiotic tolerant persister cells when challenged with either Rif/Strep or INH (Figure 1B). Rifampicin inhibits RNA polymerase, streptomycin causes mistranslation of the ribosome, and INH is a prodrug that forms and adduct with NAD, which inhibits the synthesis of mycolic acid of the mycobacterial cell wall. Tolerance of these mechanistically unrelated antibiotics shows that a decrease in ATP causes multidrug tolerance. A longer, three day treatment with bedaquiline kills *M. tuberculosis* (Koul et al., 2014), apparently lowering the concentration of ATP to a point of no return. However, the ability of bedaquiline to cause multidrug tolerance of the pathogen is a potential cause for concern and should be taken into account when developing treatment regimens.

**Fig. 1:**
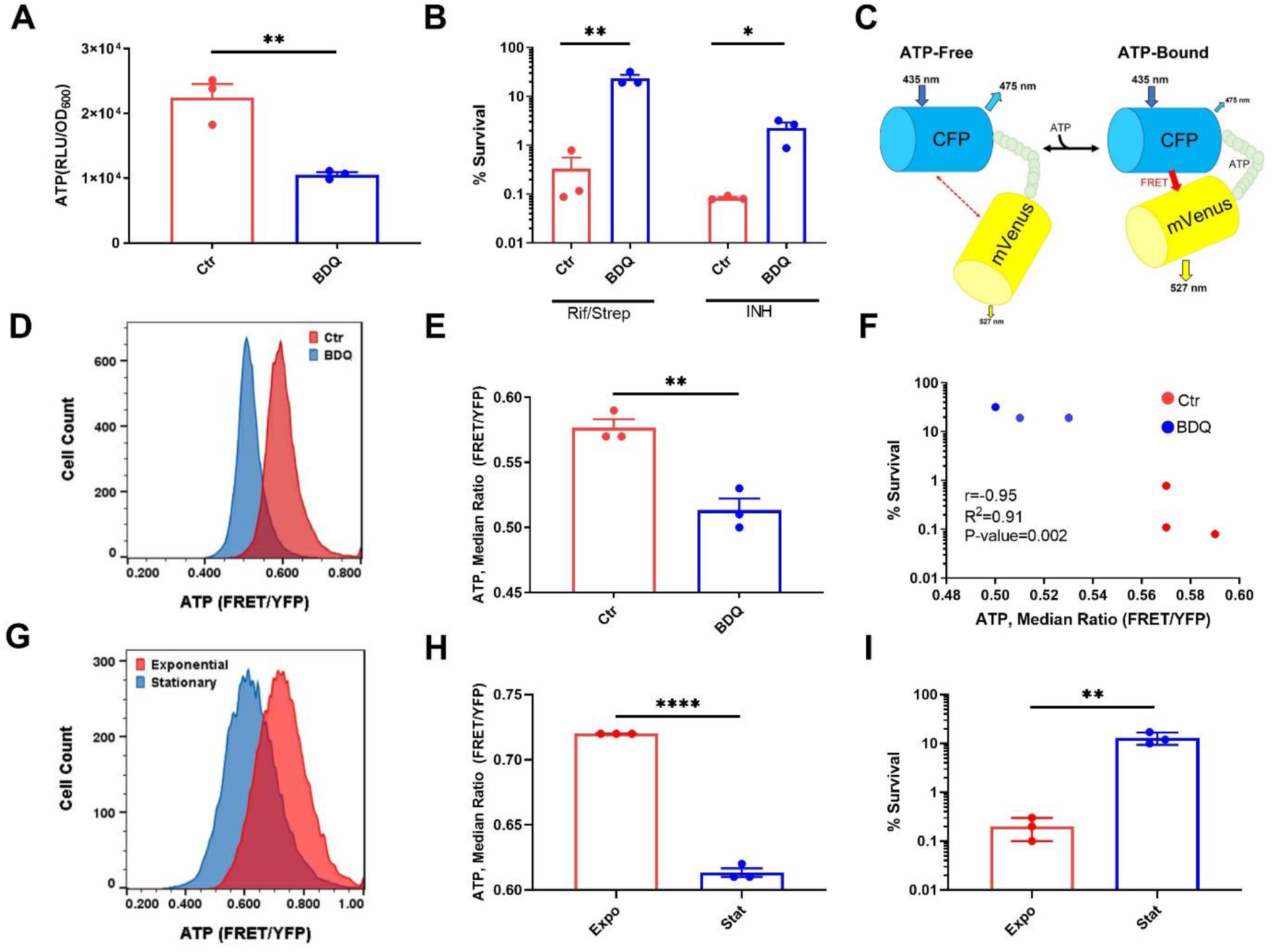
ATP level determines persister formation. Cultures of *M. tuberculosis* were grown in minimal medium with 5 mM acetate and treated with Bedaquiline (BDQ) for 4 hours (A-F) or in 7H9 rich medium (G-I). (A) Luminescence based ATP measurement after treatment with BDQ. Data are displayed as relative light units (RLU) normalized to OD_600_ of the culture. (B) Survival of BDQ treated cells after a challenge with either rifampicin (10 μg/mL) + streptomycin (10 μg/mL) or isoniazid (20 μg/mL) for 7 days. CFU/mL was determined at days 0 and 7. Data are displayed as percent survival. (C) Schematic of ratiometric FRET based ATP biosensor ATeam1.03^YEMK^. (D) Representative flow cytometry analysis of *M. tuberculosis* expressing ATeam1.03^YEMK^ treated with BDQ for 4 hours. Data displayed as FRET signal normalized to reporter expression (YFP). (E) Quantification of flow cytometry analysis in (D). Data are displayed as the median FRET/YFP ratio. (F) Correlation analysis of percent survival and median FRET/YFP ratio of untreated and BDQ treated *M. tuberculosis* cells. Data are representative of at least 3 biological replicates. Ctr=control, untreated cultures. (G) Representative example of single cell ATP analysis via ATeam1.03^YEMK^ of exponential and stationary phase *M. tuberculosis* cells. (H) Quantification of media FRET/YFP ratio in (G).(I) Survival of exponential and stationary cells after treatment with rifampicin (10 μg/mL) + streptomycin (10 μg/mL) for 7 days. CFU/mL were determined before antibiotic addition and after 7 days of treatment. P< 0.05, *, P< 0.01, **, P< 0.0001, ****. Data are representative of three biological replicates. Significance was determined by unpaired two tailed t-test.

In order to analyze ATP in single cells, we employed ATeam1.03^YEMK^, a biosensor recently developed for use in mycobacteria (Maglica, Ozdemir, & McKinney, 2015). ATeam1.03^YEMK^ is a FRET based sensor comprising a pair of cyan and yellow fluorescent proteins (CFP and YFP) flanking the epsilon subunit of the *Bacillus subtilis* F_o_F_1_ ATP synthase, which binds ATP with high affinity and specificity (Maglica et al., 2015). Binding of ATP by the epsilon subunit brings CFP in close proximity to YFP resulting in energy transfer between the fluorescent proteins. To determine ATP, FRET fluorescence is monitored using a setting optimized for excitation of CFP and emission of YFP (Figure 1C). CFP is excited at 435 nm and emission is monitored at 527 nm (YFP), CFP_ex_→YFP_em_. FRET based energy transfer from CFP to YFP is dependent on ATP concentration. The FRET values are normalized to fluorescence measurement of YFP by excitation at 488nm and collecting emission at 527nm, YFP_ex_→YFP_em_, which is not dependent on ATP concentration. This allows for normalization of cell-to-cell variation in the levels of the reporter. Normalized values are displayed as FRET/YFP and are indicative of intracellular ATP concentration. ATP was monitored in single cells of a growing culture expressing ATeam1.03^YEMK^ by FACS. Of note is the broad distribution of ATP levels among cells of the population (Figure 1D). Treatment with bedaquiline produces a distinct shift to lower levels of ATP (Figure 1D, E, F). Next, we compared ATP levels in single cells of growing and stationary cultures. As expected, ATP levels are higher in a growing population (Figure 1G, H). The level of persisters surviving treatment with Rif/Strep was 50 fold higher in the stationary population as compared to growing cells (Figure 1I), in agreement with previous findings (I. Keren et al., 2011).

We took advantage of a higher level of persisters in the stationary population to directly examine the relationship between ATP and survival in single cells using cell sorting. A gate corresponding to 2% of the population was set to sort 5,000 low or high ATP cells (Figure 2A) directly into a medium with antibiotic, and survival was monitored for 72 hours. Additionally, 5,000 cells were sorted irrespective of the ATP state of the cell (YFP only) and were considered representative of the behavior of the bulk culture. The gate was set for low ATP cells with good expression of the sensor (high YFP signal) in order to improve detection and avoid defective cells. Low ATP cells were considerably more tolerant of rifampicin (Figure 2B) or streptomycin (Figure 2C) as compared to high ATP cells or to the bulk of the population. High ATP and regular cells were essentially eliminated by rifampicin by 48 hours, while a distinct population of low ATP cells survived at 72 hours. A similar pattern was observed with a more rapidly killing streptomycin. This experiment shows that low ATP cells produced stochastically are multidrug-tolerant persisters.

**Fig. 2:**
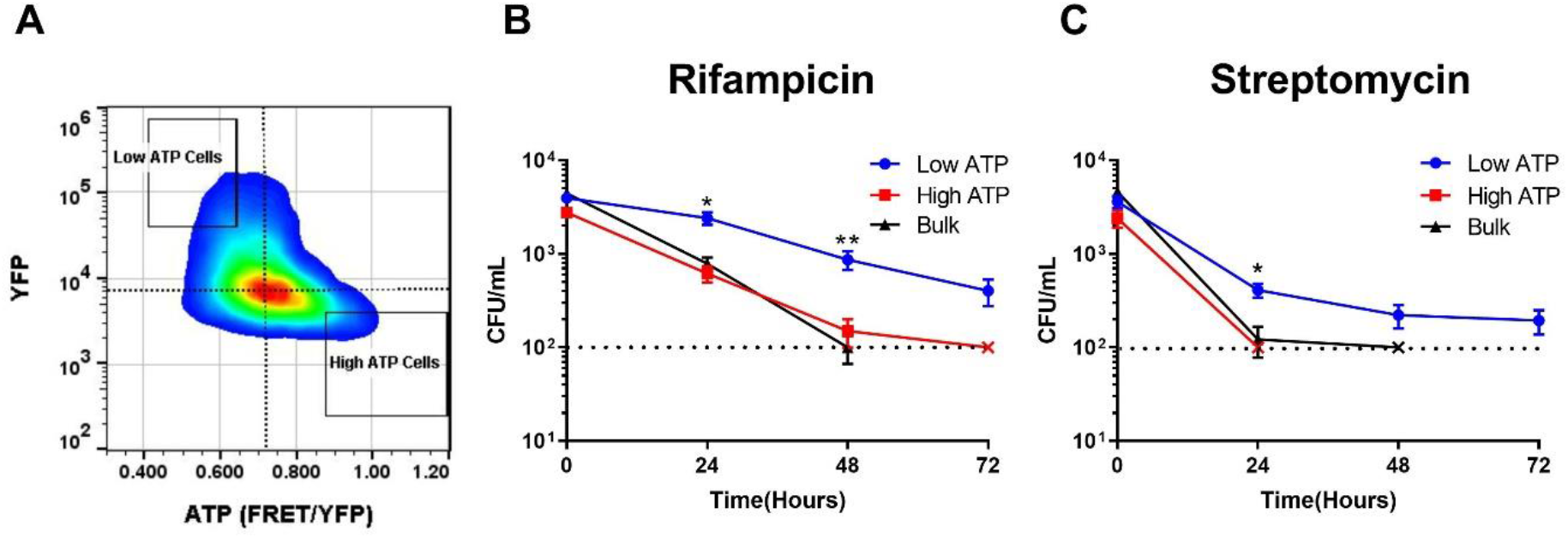
Low ATP *M. tuberculosis* cells are multi-drug tolerant. (A) Gating strategy for sorting low and high ATP *M. tuberculosis* from a population expressing ATeam1.03^YEMK^. Low and high ATP gates represent 2% of the total population. (B, C) 5,000 low, high, or bulk population cells were sorted directly into 7H9 medium containing (B) rifampicin (1 μg/mL) or (C) streptomycin (2 μg/mL). Survival was monitored by determining CFU/mL at time 0, 24, 48, and 72 hours. A dotted line represents the limit of detection. X’s indicate the time point at which a population fell below the limit of detection. P< 0.05, *, P< 0.01, **. Data are representative of at least 3 biological replicates. Significance was determined by multiple unpaired t-test.

### Identifying noise generators

Noise in the level of expression of any one among numerous enzymes participating in energy production could lead to low ATP we observe in rare single cells. In order to identify such “noisy” components, we used a simple growth medium, where the number of enzymes contributing to energy production is minimized. *M. tuberculosis* grows well in a minimal medium with acetate as a single carbon source, with a doubling time of ~21 hours, similar to its doubling time in rich media (18-20 hours). *M. tuberculosis* use two short metabolic pathways that lead from acetate to acetyl-CoA which then enters into the TCA cycle (Figure 3A). Acetate can either be converted to acetyl-CoA in a single step by acetyl-coenzyme A synthase Acs, or in a two-step pathway consisting of acetate kinase AckA producing acetyl phosphate, and the phosphotransacetylase Pta. We reasoned that consequences of an enzyme being expressed at low levels will be more apparent if the substrate is not saturating the pathway. To this end, we first tested growth of *M. tuberculosis* at different concentrations of acetate, aiming to identify a minimal level at which growth is not affected. Growth rate was similar in the presence of 20, 10 and 5 mM of acetate, and dropped at 2.5 mM acetate (Figure 3B). The level of ATP dropped in the order 20 – 10 – 5 mM acetate (Figure 3C, D), showing that growth is not affected by relatively small changes in ATP. We observed a similar trend with lactate as a single carbon source (Figure S1 A, B, C). We next analyzed the distribution variance of ATP among cells of these three populations in order to determine the relative noise in its levels (Figure 3E). For this, the coefficient of variation (CV) was derived from the FRET/YFP distributions generated via single cell FACS analysis (Figure 3C). CV quantifies variance by dividing the standard deviation (σ) of FRET/YFP ratio of cells in a population by mean (μ) FRET/YFP ratio of the population (σ/μ). Noise in ATP levels among cells decreased in the order 5 – 10 – 20 mM acetate (Figure 3E, F). Again, we observed a similar phenotype in a medium with lactate (Figure S1D, E). This suggested that more persisters will form in a medium with lower acetate or lactate. Indeed, a population growing in 5 mM acetate, or 10mM lactate, produced about 10 fold more persisters as compared to cells in a 20 mM sample (Figure 3G, Figure S1F). Survival is negatively correlated with median ATP of the population (Figure 3H) and positively correlated with noise in ATP level (Figure 3I). Notably, apart from linking noise to persister formation, this experiment shows that growth rate per se does not determine tolerance.

**Fig. 3:**
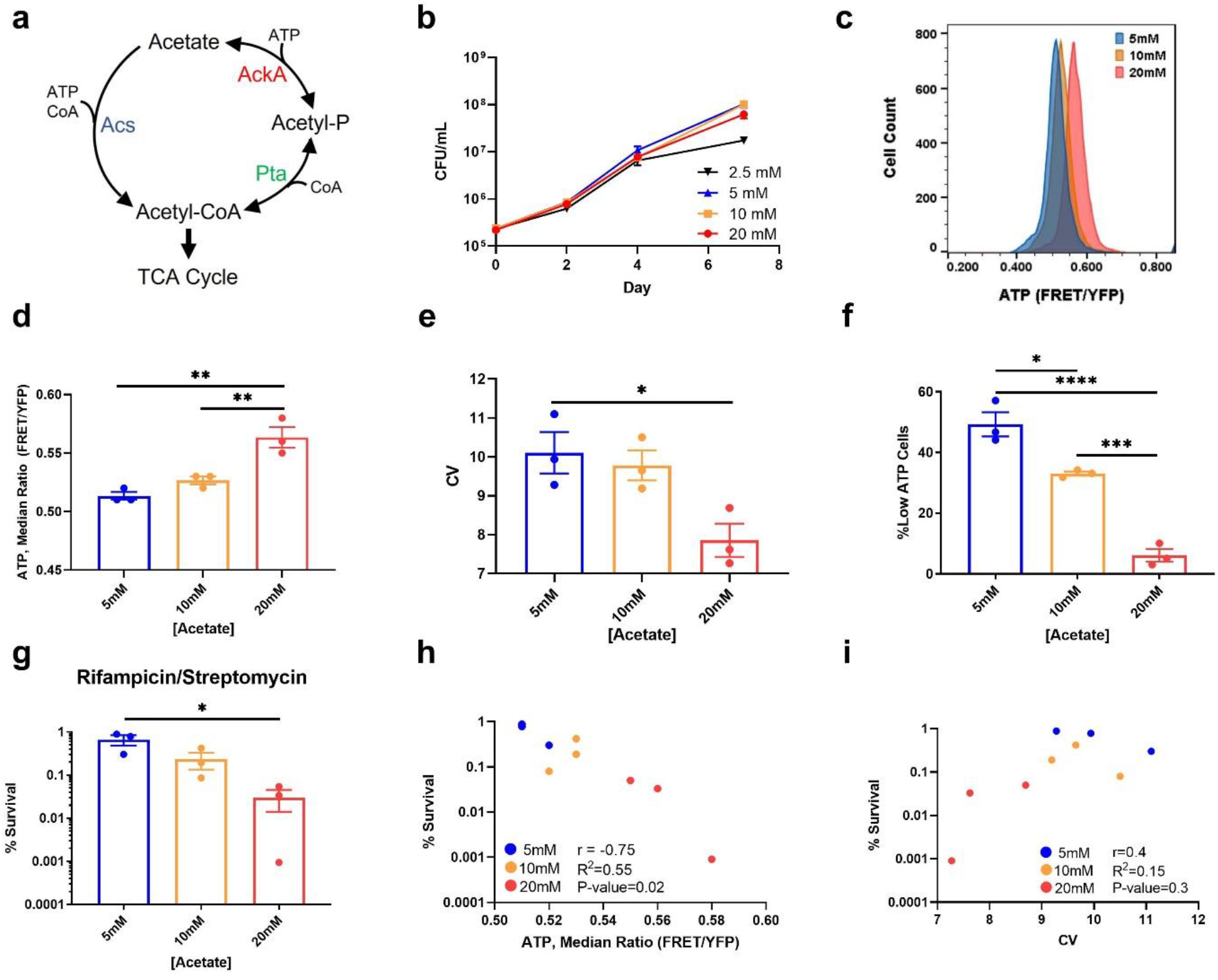
Limiting acetate increases noise in ATP levels, and persisters. (A) Schematic of acetate catabolism genes in *M. tuberculosis*. Acetate is converted to acetyl- CoA by Acs in a single step reaction or by AckA and Pta in a two-step reaction.(B) Growth curve of *M. tuberculosis* in minimal media with varying concentrations of acetate as the sole carbon source. (C) Representative example of flow cytometry analysis of *M. tuberculosis* expressing ATeam1.03^YEMK^. *M. tuberculosis* was grown in minimal media with the indicated concentrations of acetate for 1 week before being analyzed. (D) Quantification of median FRET/YFP ratio generated by Ateam1.03^YEMK^ in (C). **e,** Quantification of coefficient of variation (CV) of FRET/YFP ratio in (C). **f,** Quantification of “Low ATP Cells” defined as events falling below a gate set at FRET/YFP ratio one standard deviation below median ratio in 20 mM sample, the sample with the highest median FRET/YFP ratio. (G) Survival of *M. tuberculosis* grown at indicated concentrations of acetate after being challenged with rifampicin (10 μg/mL) + streptomycin (10 μg/mL) for seven days. (H) Correlation analysis of survival and median FRET/YFP ratio of populations. (I) Correlation analysis of survival and CV of FRET/YFP ratio. P< 0.05, *, P< 0.01, **, P< 0.001, ***, P< 0.0001, **** Data are representative of at least three biological replicates. Significance determined by one-way anova with Tukey’s post test.

We then used single cell reverse transcription quantitative PCR (RT-qPCR) recently developed for mycobacteria (Srinivas et al., 2020) to directly link noise in ATP concentration to the level of expression of enzymes in the acetate metabolic pathways. For this, we used cells growing at a sub-optimal level of acetate (2.5 mM) in order to maximize possible cell-to-cell differences in ATP. Twelve individual low and high ATP cells were sorted into a microtiter plate and transcripts of the acetate metabolism genes *acs, ackA*, and *pta* were measured by RT-qPCR. The low ATP cells had lower levels of transcripts, while the high ATP cells produced more transcripts (Figure 4A, each dot represents measurement from a single cell). There was a particularly wide variation in the expression levels of *ackA* coding for acetate kinase.

**Fig. 4:**
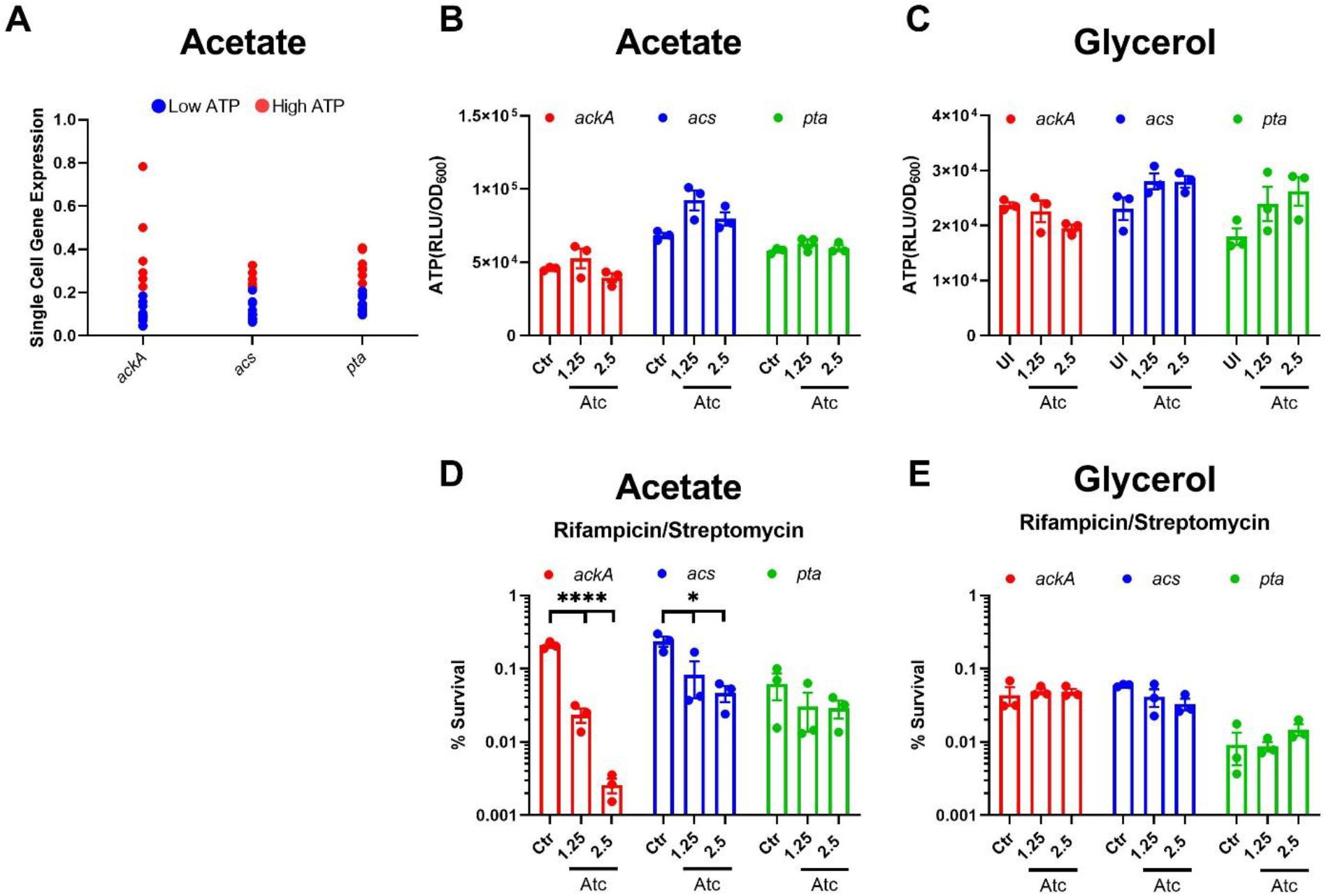
Noise quenching in *ackA* expression reduces drug tolerant persisters. (A) Single cell expression analysis of low and high ATP cells sorted from a minimal medium with 2.5 mM acetate as the carbon source. *M. tuberculosis* expressing ATeam1.03^YEMK^ was analyzed via FACS and a single low or high ATP cell was dispensed into a well of a 96 well qPCR plate. “Low” and “High” ATP cells were gated as in Figure 2A. Normalized expression of the indicated genes was determined. Expression was normalized to the threshold cycle (Ct) value determined from the origin of replication of pND235-YEMK (plasmid expressing ATeam1.03^YEMK^). (B,C) ATP in a population of *M. tuberculosis* overexpressing *acs, ackA*, or *pta* in minimal media with 2.5mM acetate (B) or 0.01% glycerol (C) as the sole carbon source. ATP was measured by luciferase after one week of growth, with or without induction with anhydrotetracycline (Atc). Luminescence is normalized to the OD_600_ of the culture. (D, E) Survival of *M. tuberculosis* expressing *acs, ackA*, or *pta* under control of a tetracycline inducible promoter. Cultures were grown in minimal media with (D) 2.5 mM acetate or (E) 0.01% glycerol as the sole carbon source for 7 days. The cultures were left uninduced (Ctr) or induced with Atc. Cultures were then challenged with rifampicin (10 μg/mL) + streptomycin (10 μg/mL) for 7 days. CFU/mL were determined before antibiotic treatment and after 7 days of treatment. P< 0.05, *, P< 0.0001, ****. Data are representative of at least three biological replicates. Significance was determined by one-way anova with Tukey’s post test.

We reasoned that quenching the noise will diminish persisters if it is indeed responsible for their formation. Ectopic overexpression of an enzyme should quench the noise in its expression. To test this, we cloned *ackA, acs* and *pta* genes into a chromosomally integrating plasmid under the control of a tetracycline inducible promoter. Cultures were grown in minimal media with acetate in the presence of anhydrotetracycline (Atc). Overexpression did not affect growth of these strains (Figure S2). Regardless of carbon source, no change in ATP was seen in the bulk population induced with Atc (Figure 4B, C). This is to be expected, since only persisters that express lower amounts of energy producing components will be sensitive to a decrease in substrate concentration. Changes in the metabolic state of this small subpopulation will not affect bulk ATP measurement. Strikingly, induction of *ackA* resulted in a 100 fold decrease in persisters tolerant of killing by Rif/Strep (Figure 4D). There was a notable but considerably smaller decrease in persisters upon overexpression of *acs*, and no change in cells overexpressing *pta*. Importantly, survival was unaffected when these genes were overexpressed in minimal media with suboptimal glycerol (0.01%, Figure S3) as the sole carbon source (Figure 4E). These results suggest that *ackA* is the main noise generator in the acetate metabolic pathway and acts as a persister gene.

## Discussion

Studies of antibiotic tolerance in *M. tuberculosis* and persister cells in other bacterial species have proceeded along parallel lines, with little cross-talk. There is a considerable body of evidence showing that metabolic downshift by a variety of means and mechanisms leads to antibiotic tolerance in different bacterial species (Amato, Orman, & Brynildsen, 2013; Shatalin et al., 2021). Of particular relevance to tuberculosis is the finding that establishing dormancy in *M. tuberculosis* under hypoxia involves a metabolic switch to synthesis of triglycerides, diminishing the available energy resources for other biosynthetic functions, and contributing to antibiotic tolerance (Baek et al., 2011). Our recent work established that both *E.coli* and *S. aureus* persisters are low ATP cells (Brian P. Conlon et al., 2016; Shan et al., 2017). Consistent with this, we show that *M. tuberculosis* persisters are also low ATP cells (Figure 1 and 2). Additionally, as with *E. coli* persisters (Manuse et al., 2021), *M. tuberculosis* persisters are stochastically formed during normal growth (Figure 2). These data suggest, as with other bacteria, that a low energy state underlies persister formation in *M. tuberculosis*.

In *S. aureus* and *E. coli*, sorting of cells with low expression of Krebs cycle enzymes enriches in persisters (Brian P. Conlon et al., 2016; Shan et al., 2017; Zalis et al., 2019). This suggests that noise in expression of energy generating pathways drives a low ATP state and persister formation. To examine this relationship more directly we made use of simple medium with acetate as the sole carbon source. We reasoned that when substrate is limiting, noise in energy generating pathways will become more pronounced and lead to increases in low ATP cells and persisters. At a minimal concentration of acetate that does not yet diminish growth, ATP level drops with a concomitant increase in persisters (Figure 3 and S1). To monitor ATP and noise in energy-generating component in the same cell, we employed single cell transcription analysis of sorted cells with low ATP determined by a specific fluorescent reporter ATeam1.03^YEMK^. This shows considerable noise in the expression of the acetate kinase AckA in low ATP *M. tuberculosis* cells when acetate is the sole carbon source (Figure 4A). Further, by quenching noise through overexpression of AckA we were able to dramatically decrease persisters (Figure 4D) identifying AckA as the principal noise generator under these conditions.

Apart from a general low-energy mechanism, several specialized mechanisms have been described in bacteria that operate under particular conditions. In *E. coli*, DNA damage by fluoroquinolone antibiotics induces the SOS response leading to expression of the TisB toxin that forms an ion channel in the membrane, decreases the pmf and ATP, and is primarily responsible for persister formation under these conditions (Berghoff, Hoekzema, Aulbach, & Wagner, 2017; Dorr, Vulic, & Lewis, 2010; Gurnev, Ortenberg, Dorr, Lewis, & Bezrukov, 2012). In *E. coli*, a gain of function hipA7 (high persister) mutation in the HipA toxin produces hip mutants (Kaspy et al., 2013; Moyed & Bertrand, 1983) that are found in patients treated for urinary tract infection (Schumacher et al., 2015). Whether specialized mechanisms of persister formation exist in *M. tuberculosis* is an important open question. This is especially significant since mutations leading to increased tolerance have been shown to favor development of classical resistance in *E. coli* and *S. aureus* (Balaban, Merrin, Chait, Kowalik, & Leibler, 2004; Levin-Reisman et al., 2017; J. Liu, Gefen, Ronin, Bar-Meir, & Balaban, 2020; Moreno-Gamez et al., 2020). In tuberculosis, failure to clear the infection due to tolerance has been linked to development of resistance but whether hip mutations in *M. tuberculosis* favor selection of resistant mutants remains to be established.

A number of studies have linked persisters to disease, starting with the isolation of *P. aeruginosa* hip mutants from patients with cystic fibrosis undergoing lengthy antibiotic therapy (Bartell et al., 2021; Mulcahy, Burns, Lory, & Lewis, 2010). *hip* mutants have been identified in clinical isolates of *M. tuberculosis* as well (Torrey et al., 2016). In the case of *Salmonella*, entrance of cells into macrophages results in a dramatic increase in persisters (Helaine et al., 2014). *M. tuberculosis* similarly colonizes macrophages, upon which antibiotic tolerance of the bulk population increases (Y. Liu et al., 2016; Pieters, 2008). In *S. aureus*, a decrease in ATP and antibiotic tolerance in vivo can result from inhibition of respiration by compounds originating from a co-infection with *P. aeruginosa*, and by ROS produced by macrophages (Huemer et al., 2021; Radlinski et al., 2017; Rowe et al., 2020).

In *M. tuberculosis*, there are two different paths leading to quiescence, and both are likely responsible for the lengthy antibiotic therapy required to treat tuberculosis. A population entering into dormancy in response to external stressors such as hypoxia, and stochastically formed persisters appear to share the same basic mechanism of antibiotic tolerance – a low energy state. This suggests that a therapeutic approach against dormant cells will be effective irrespective of which path they used to enter into dormancy. However, while we have a sizable and growing (albeit slowly) arsenal of antibiotics that act against regular cells, discovery of anti-persister compounds is still in its infancy (Lewis, 2020), with but a few examples of anti-persister compounds(Brotz-Oesterhelt et al., 2005; B. P. Conlon et al., 2013; Griffith et al., 2019; Kim et al., 2018). An alternative approach is pulse-dosing with conventional antibiotics that has been described for eradicating a biofilm formed by *S. aureus* in vitro (Meyer, Taylor, Seidel, Gates, & Lewis, 2020). Combatting dormancy will require novel types of compounds and approaches.

Our findings suggest a general model for persister formation in bacteria, which is shared by *M. tuberculosis* (Figure 5). Noise in the expression of an energy-generating component such as *ackA* results in rare cells that have low levels of ATP. This in turn will decrease the activity of targets, preventing antibiotics from corrupting them. The nature of the principal noise generator will depend on which metabolic pathway is dominant under given growth conditions. From this perspective, there will be many “persister genes” in *M. tuberculosis* and other bacteria. Noise quenching that we describe in this study provides a direct means to test the involvement of a given gene in persister formation. This approach should also be applicable *in vivo*, where tetracycline inducible gene expression has been used. Notably, noise quenching by overexpression provides a simple approach to quantitatively determine the relative input of any gene into persister formation.

**Fig. 5:**
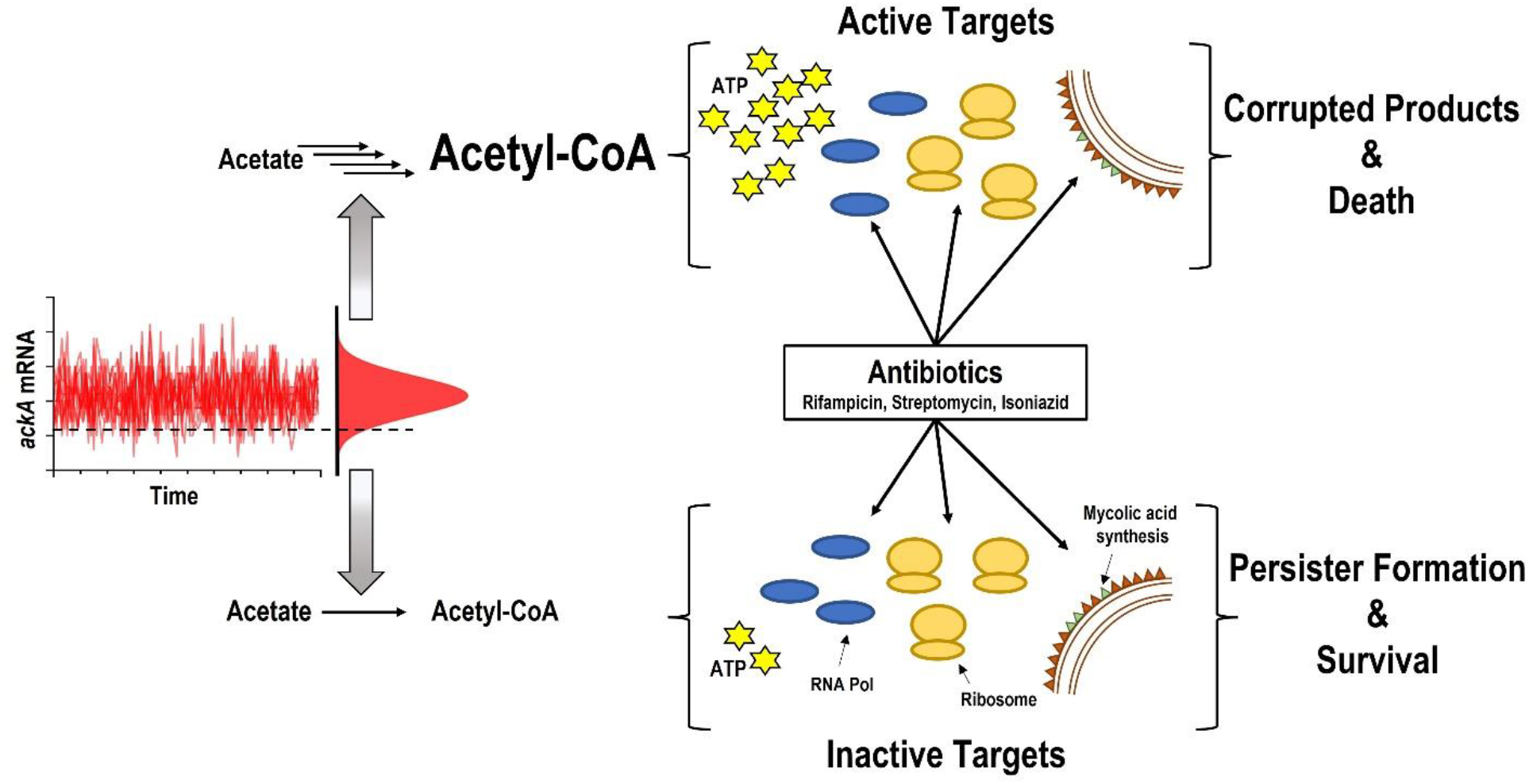
A model of persister cell formation. When *M. tuberculosis* is grown in minimal media with acetate, the acetate kinase AckA represents a bottleneck in the energy-producing pathway. In the majority of cells, AckA is not limiting and allows for the efficient catabolism of acetate. This translates into higher levels of ATP, and active targets such as RNA polymerase, the ribosome, and mycolic acid synthesis. Antibiotics corrupt the targets resulting in cell death. Noise in transcription causes stochastic decreases in AckA. This leads to a decrease in ATP and inactive targets, creating a multidrug-tolerant persister cell.

## Materials and Methods

### Bacterial strains and media

*M. tuberculosis* strain used for all experiments was H37Rv MC^2^6020. *M. tuberculosis* was grown in Difco 7H9 supplemented with 10% OADC, 5% glycerol, lysine (80 μg/mL), pantothenate (24 μg/mL), and 0.05% tyloxapol. For CFU enumeration *M. tuberculosis* was plated on Difco 7H10 supplemented with 10% OADC, 5% glycerol, lysine (80 μg/mL), and pantothenate (24 μg/mL). To make minimal media 0.5 g asparagine, 1 g NH2PO4, 2.5 g Na2HPO4, 50 mg ferric ammonium citrate, 0.5 g MgSO4, 0.5 mg CaCl2, and 0.1 mg ZnSO4 were dissolved in 1 L of water. Lysine (80 μg/mL), pantothenate (24 μg/mL), and tyloxapol (0.05%) were added and the media was filter sterilized. Carbon sources were added to the media at the indicated concentrations prior to experiment. For construction of overexpression strains, genes were amplified using primers in supplementary table xx and cloned into plasmid pTetSGkan. Plasmids were transformed into *M. tuberculosis* and selected on 7H10 complete media supplemented with 40 μg/mL Kanamycin.

### Growth measurements

For analysis of growth in minimal media with single carbon sources, *M. tuberculosis* were grown in 7H9 complete media, washed twice in PBS, then resuspended in minimal media with the indicated carbon source to an OD600 ~0.01 at a volume of 10mL. The cultures were grown for a week with shaking at 37C. Time points were plated for CFU at Day 0, 2, 4, and 7.

### Antibiotic survival assay

For analysis of bedaquiline effects on survival, cultures were challenged in exponential phase when OD600 ~0.8. For analysis of survival in exponential and stationary phase, cultures were challenged when OD600 ~0.8 (exponential) or ~1.8-2 (Stationary). For analysis of survival in minimal media as well as overexpression analysis in minimal media, cultures were grown for 7 days under the indicated conditions and then challenged. In all cases, an aliquot was removed before treatment, serial diluted, and plated for day 0 CFU enumeration. Cultures were treated with the indicated antibiotic(s) for 7 days after which an aliquot was removed, washed once in PBS, serial diluted and plated for CFU. Percent survival was calculated as follows: (CFU Day 7/ CFU Day 0)*100. For analysis of survival in bulk cultures, antibiotic concentrations used were Rifampicin (10 μg/mL) + Streptomycin (10 μg/mL) or Isoniazid (20 μg/mL). Sorted cells were treated with Rifampicin (1 μg/mL) or Streptomycin (2 μg/mL).

### ATP Quantification

Prior to antibiotic treatment aliquots of cultures were washed with PBS, and intracellular ATP concentration was measured by BacTiter Glo kit (Promega, Madison, WI, USA) according to the manufacturer’s instructions. Bioluminescence values (RLU) were normalized to OD_600_.

### Flow Cytometry and Fluorescent Activated Cell Sorting

Single cell ATP was analyzed using *M. tuberculosis* expressing pND235-YEMK. This plasmid encodes a FRET based ATP biosensor adapted for use in *M. tuberculosis* (Maglica et. al., 2015). FRET-based fluorescence of single cells were collected on BD FACS Aria II flow cytometer (BD Biosciences, San Jose, CA, USA) with a 70-μm nozzle. Fluorescence was collected for YFP emission at two separate laser excitations with band pass filters optimized for YFP, excitation at 445nm (CFP_ex_ → YFP_em_) (FRET) and excitation at 488nm (YFP_ex_ → YFP_em_). Single cell normalized ATP is expressed as the FRET/YFP ratio [(CFP_ex_ → YFP_em_)/(YFP_ex_ → YFP_em_)]. For flow cytometry analysis of cultures expressing pND235-YEMK, a minimum of 20, 000 events were collected. The events were gated for size (FSC-A, SSC-A), YFP positivity, and finally ratiometric signal of ATP (FRET/YFP) was determined. Aliquots of the cultures were directly analyzed on the flow cytometer in the experimental media conditions. All analysis was conducted in FlowJo (BD). For survival sorting experiments, a liquid culture of *M. tuberculosis* expressing pND235-YEMK was grown to stationary phase and then diluted 1:100 in fresh 7H9 media. The culture was grown to late stationary phase (~2 weeks), diluted 1:20 in PBS, and loaded onto the BD FACS Aria II. To sort based on the normalized ratiometric signal from pND235-YEMK, the ratio feature of the FACS Diva software was enabled to calculate FRET/YFP signal in real time. Events were first gated for size (FSC-A, SSC-A), and YFP positivity, followed by analysis of single cells as a dot plot of FRET/YFP versus YFP signal. “Low ATP” and “High ATP” gates were set to 2% of the total population. YFP+ cells were gated as any cell expressing YFP above background levels. A total of 5,000 low ATP, high ATP, or YFP+ cells were sorted directly into 1 mL 7H9 liquid media containing either rifampicin (1 μg/mL) or streptomycin (2 μg/mL). The sample was immediately serial diluted and plated on 7H10 to determine Day 0 CFU/mL. The sample was then plated to determine CFU/mL on days 1, 2, and 3 post-treatment. At all timepoints, a 95 μL aliquot of each sample was removed and added to 5 μL 1% activated charcoal (final concentration of activated charcoal is 0.05%) before being serial diluted to limit the effects of the antibiotics in the media on CFU determination. Two technical replicates were collected per experiment reversing the order of sample collection between replicates to limit effects of collection timing bias. The experiment was repeated a minimum of three times.

### Single cell RT-qPCR

Protocol for single cell RT-qPCR was based on a recently developed method (Srinivas et. al., 2020). Cultures expressing pND235-YEMK growing in minimal media with 2.5 mM acetate as the sole carbon source were grown to stationary phase. Low ATP, High ATP, and YFP+ cells were sorted based on the criteria described above. Single Low ATP, High ATP, or YFP+ cells were sorted directly into individual wells of a 96 well Bio-Rad (Hercules, CA, USA) PCR plate containing 2 μL lysis solution comprising 10% NP-40, SuperScript IV Vilo master mix (Invitrogen, Waltham, MA, USA), T4 Gene32 (Thermofisher, Waltham, MA, USA), SUPERase RNase Inhibitor (Invitrogen), and 10 pM RNA spike in control. Firefly luciferase (FLuc) RNA served as the spike in control and was generated using HiScribe T7 Quick RNA Synthesis kit (NEB, Ipswich, MA, USA) using linearized plasmid DNA encoding FLuc gene as template. A total of 16 Low ATP, High ATP, and YFP+ cells were sorted per experiment. After sorting, the cells were lysed by first flash freezing in liquid nitrogen, then placing the plate at −80C for 1 hour, followed by thawing at room temperature. The plate was then transferred to a thermocycler and cDNA was generated following cycling conditions in manufacturers protocol. Following cDNA synthesis, 25 cycles of pre-amplification was conducted using gene specific primers. Excess primer was then removed via incubation with Exonuclease I (Thermofisher) for 1 hour at 37C. One tenth (2 μL) of pre-amplified cDNA was then used to assess the single cell expression of each gene of interest using BioRad SsoAdvanced Universal SYBR Green Supermix on a BioRad CFX96 system. Amplification of the E. coli origin of replication (oriE) from pND235-YEMK served as the lysis control. Wells that generated no amplification or a threshold cycle (Ct) > 40 for oriE were considered failed lysis and removed from analysis. For all other genes, amplification values with Ct > 40 were removed from analysis. Ct values of genes were corrected for the difference between FLuc Ct in each well with the median FLuc Ct. Finally, expression of each gene was normalized to oriE amplification in each well as it is assumed oriE exist as a single copy. All primers used for single cell RT-qPCR analysis can be found in Table S1.

### Statistics

All statistical analysis was conducted using GraphPad Prism V 9.

## Acknowledgements

This work was supported by grant R01 AI141966 from the NIH to K.L. Plasmid pND235-YEMK was a kind gift from John D. McKinney at the School of Life Sciences, Swiss Federal Institute of Technology in Lausanne (EPFL), Lausanne, Switzerland.

---

**Figure S1:**
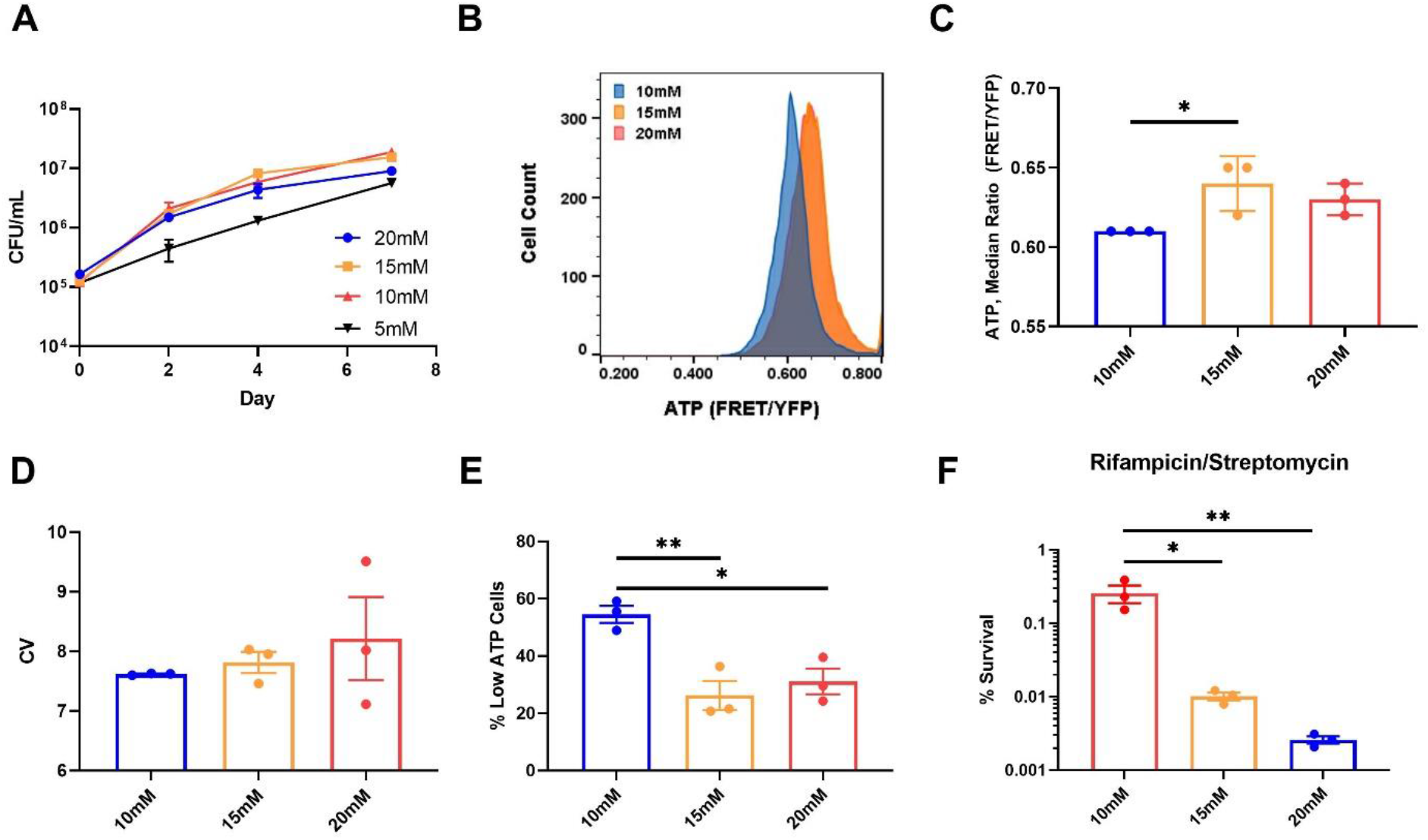
Limiting lactate increases noise in ATP levels, and persisters. (A) Growth curve of *M. tuberculosis* in minimal media with varying concentrations of lactate as the sole carbon source. (B) Representative example of flow cytometry analysis of *M. tuberculosis* expressing ATeam1.03^YEMK^. *M. tuberculosis* was grown in minimal media with the indicated concentrations of lactate for one week before being analyzed. (C) Quantification of median FRET/YFP ratio generated by ATeam1.03^YEMK^ in (B). (D) Quantification of co-efficient of variation (CV) of FRET/YFP ratio signal in (B). (E) Quantification of “Low ATP Cells” defined as events falling below a gate set at FRET/YFP ratio one standard deviation below median ratio in 20 mM sample, the sample with the highest median FRET/YFP ratio. (F) Survival of *M. tuberculosis* grown at indicated concentrations of lactate after being challenged with rifampicin (10 μg/mL) + streptomycin (10 μg/mL) for seven days. P< 0.05, *, P< 0.01, **. Data are representative of at least three biological replicates. Significance was determined by one-way anova with Tukey’s post test.

**Figure S2:**
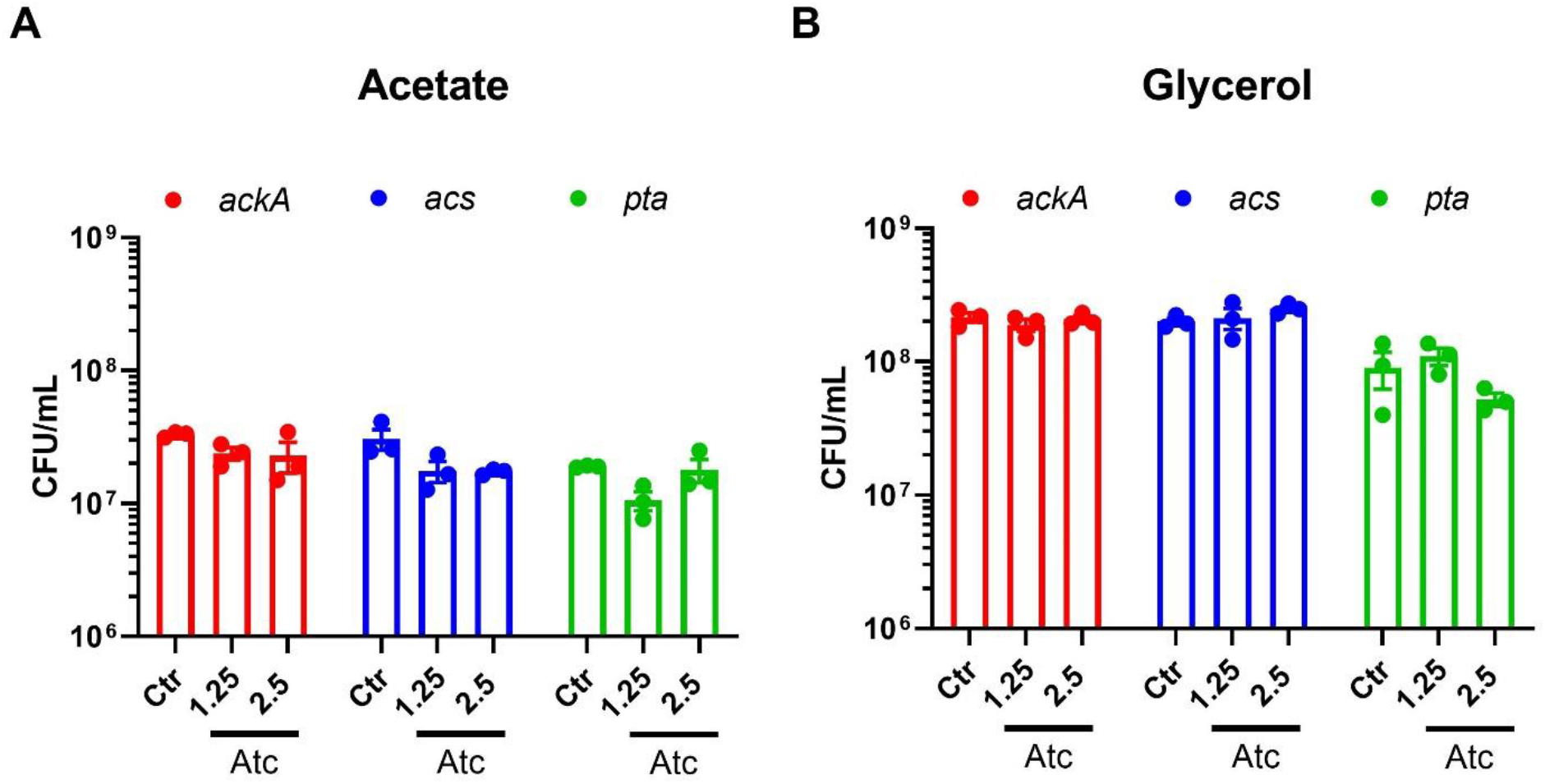
Overexpression does not affect growth of *M. tuberculosis*. Initial CFU/mL of *M. tuberculosis* expressing *acs, ackA*, or *pta* under control of a tetracycline inducible promoter. Cultures were grown in minimal media with (A) 2.5 mM acetate or (B) 0.01% glycerol as the sole carbon source for 7 days. The cultures were left uninduced (Ctr) or induced with Atc. Data are representative of three biological replicates.

**Figure S3:**
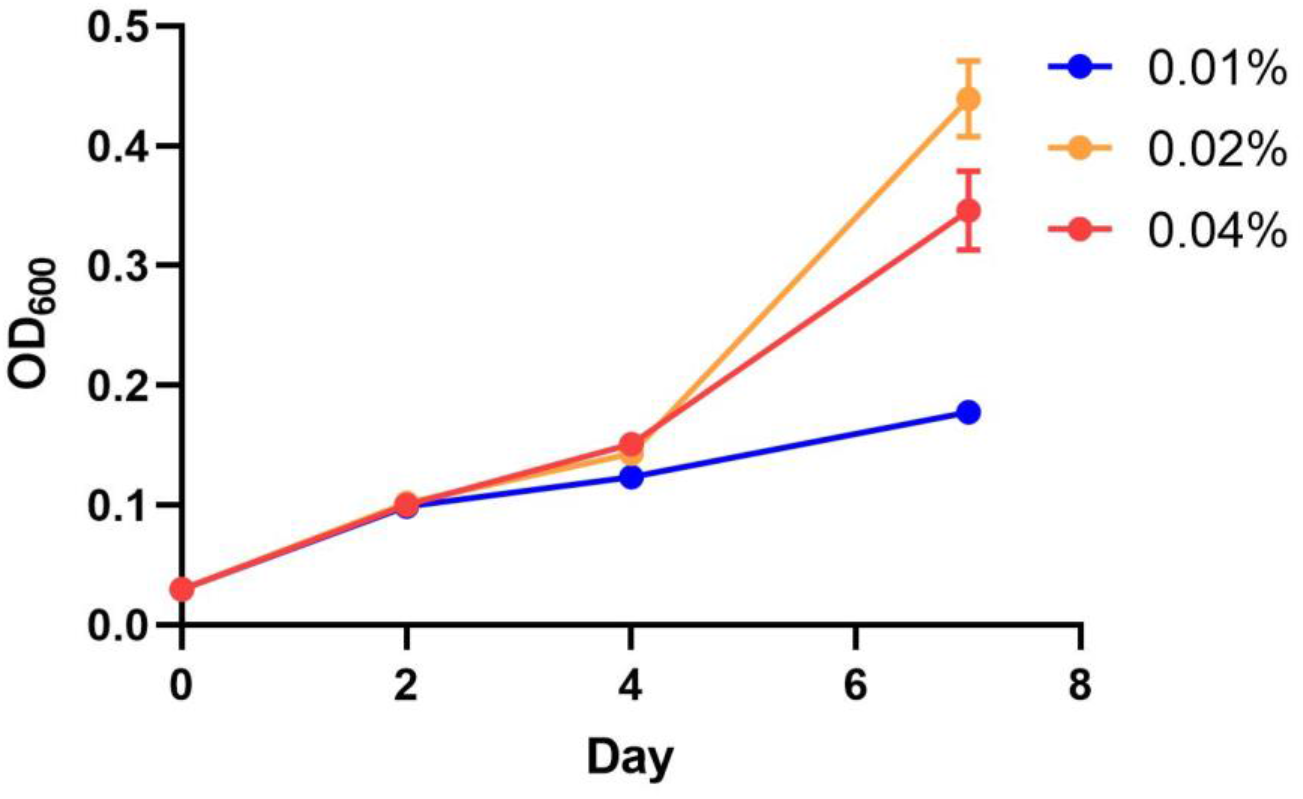
Growth curve of *M. tuberculosis* in glycerol. Growth of *M. tuberculosis* in minimal media with carrying concentrations of glycerol as the sole carbon source. Data are representative of three biological replicates.

**Table S1:**
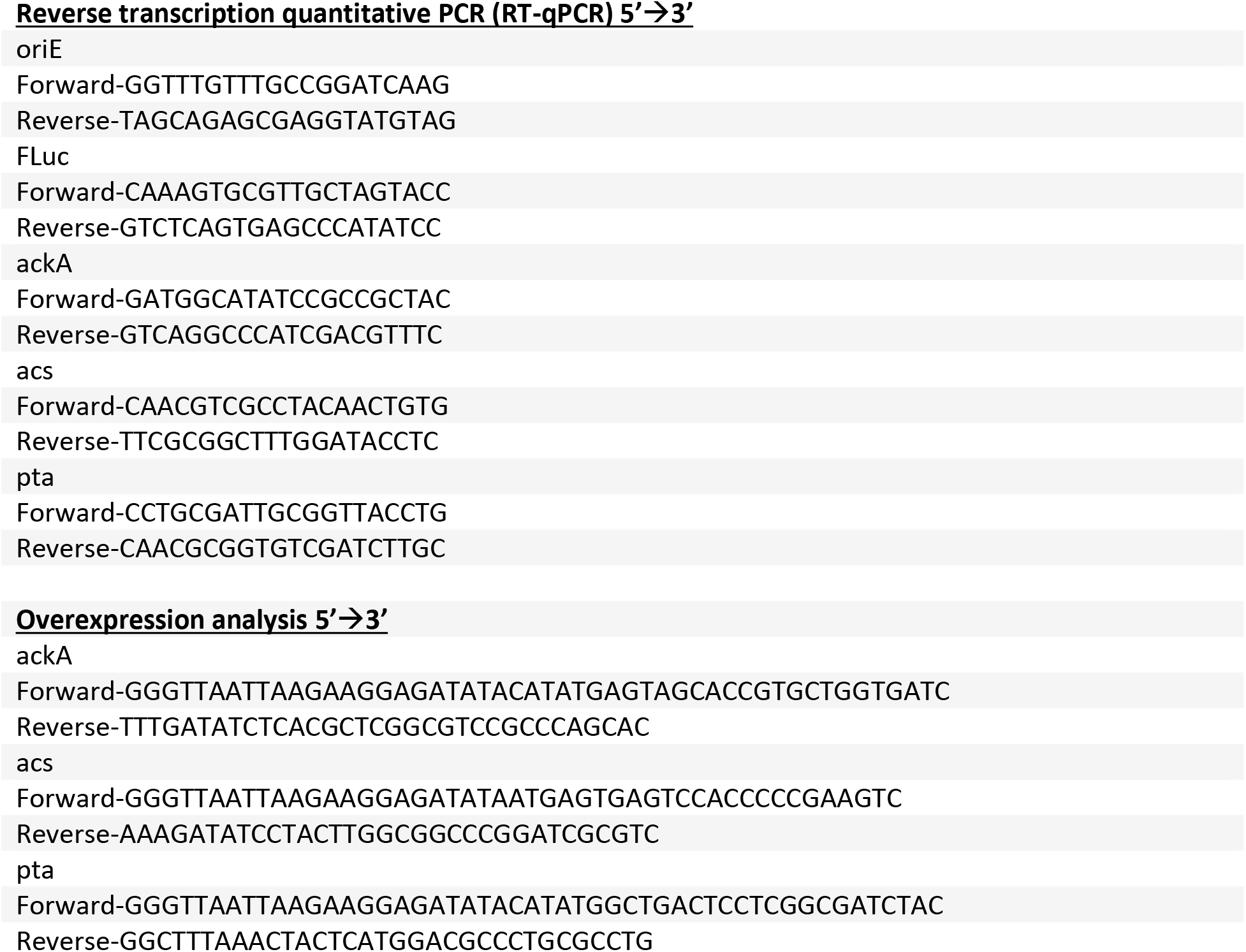
Primers used in this study.

